# Novel pathways converge with quorum sensing to regulate plant and insect host-specific factors in *Erwinia carotovora*

**DOI:** 10.1101/2023.03.13.532345

**Authors:** Filipe J. D. Vieira, Luís Teixeira, Karina B. Xavier

## Abstract

*Erwinia carotovora Ecc15* is a vector-borne phytopathogen that relies on insects to be transmitted between plant hosts. To interact with its hosts, this bacterium depends on host-specific bacterial traits. Plant tissue maceration depends on production of plant cell wall degrading enzymes (PCWDE), while survival in the digestive tract of the insect requires the *Erwinia* virulence factor (*evf*). Evf expression is responsible for the cost of *Ecc15* infection in *Drosophila melanogaster* and overexpression is lethal to the insect host. Therefore, its expression must be well controlled. Expression of *evf* and PCWDEs is co-regulated by quorum sensing via the transcriptional regulator Hor. Since virulence factors are often controlled by multiple signals, we asked which additional factors regulate *evf* expression. Using a genetic screen, we identified the sensor histidine kinase *arcB* and a new TetR-like regulator (named herein as *lvtR*, after **L**ow **V**irulence **T**ranscriptional **R**epressor), as novel regulators not only of *evf*, but also of *pelA*, which encodes a major PCWDE. We further demonstrate that *arcB* and *lvtR* mutants have reduced plant tissue maceration and reduced development delay and lethality in *Drosophila melanogaster*, compared to wild-type bacteria. Thus showing the importance of these regulators in the establishment of *Erwinia*-host-vector interactions. We also found that ArcB and LvtR regulation converges on Hor, independently of quorum sensing, to co-regulate expression of both plant and insect bacterial interaction factors during plant infection. Taken together, our results reveal a novel regulatory hub that enables *Ecc15* to integrate quorum sensing responses and environmental cues to co-regulate traits required for infection of both the plant and the insect vector. Moreover, we show that ArcB regulation of bacteria-host interaction processes is conserved in other bacteria.

**Author Summary:** Vector-borne pathogens depend on continuous cycles of replication and transmission between hosts and vectors, requiring multiple factors to interact with each of the hosts. The expression of these multiple interaction factors can be very costly, so it is expected that regulation of virulence has been evolutionarily tuned to produce expressions patterns that minimize the cost of establishing the infection while maximizing transmission efficiency of the pathogen. Here, we investigate the tripartite interaction between *Ecc15*, a plant and an insect, and show that quorum sensing, a widely conserved sensory regulator *arcB* and a regulator of previously unknown function, *lvtR*, converge to simultaneously co-regulate the expression of bacterial factors required for these interactions. Gene expression regulation is channeled through the conserved regulator Hor, which serves as a molecular hub for the integration of these multiple signals. Our data suggest that integration of multiple signals to co-regulate plant and insect associated factors ensure fine-tune titration of gene expression and maximization of bacterial energetic resources.

## Introduction

Vector-borne phytopathogens often rely on insects to be transmitted between plant hosts. These insects by feeding on plant sap or in rotting tissues, caused by the pathogen itself, can acquire the pathogen, and subsequently transmit it (1, 2). The cell and molecular biology of plants and insects is substantially different, therefore the traits required for the interaction between the host or the vector are likely different, and specific to each host (3). When bacterial genes are linked to pathology (manifestation of disease) and/or cause damage to the host (tissue rotting, as an extreme case), they are typically classified as virulence factors (4). To ensure a continuous transmission cycle, a fine balance between production of virulence factors and their fitness cost for the microbe, vector and host is necessary (5, 6). To achieve this balance, tight control of expression of virulence factors is required (7–9). Identification of the molecular mechanisms and signaling networks involved in regulation of virulence expression is thus crucial for understanding the maintenance of the transmission-infection cycle, the establishment of host-pathogen-vector interactions and, importantly, for designing strategies aiming at interfering with such processes.

*Erwinia carotovora* is a plant pathogen that causes soft-rot disease in many economically relevant crops (10). These phytopathogens can enter the plant vascular system either by natural plant openings, like the stomata, or by wounds caused by herbivorous insects (2, 10). Once inside the plant host, these bacteria disrupt the normal functioning of cells by producing a battery of virulence factors: the plant cell wall degrading enzymes (PCWDE). PCWDE are regulated in a cell density dependent manner via quorum sensing (11–13), and also by environmental cues, such as 2-keto-3-deoxygluconate (KDG), 2,5-diketo-3-deoxygluconate or polygalacturonate, intermediate molecules that are formed after pectin degradation (14–16). By targeting the cell wall, PCWDEs compromise its integrity, leading to cell lysis, and subsequent tissue decay (10, 17).

*Erwinia* have also been used to study the mechanisms of association with insects. Importantly, *Erwinia* survives poorly in soil, requiring insects to continue its infection cycle (10). The strain *Ecc15* has been used to study the molecular mechanisms that regulate the interaction between *Erwinia carotovora* and *Drosophila melanogaster* (18). In this strain, the gene *evf* (*Erwinia* virulence factor) promotes the survival of *Ecc15* in the gut of *Drosophila melanogaster* (19, 20). However, the increase in the survival of the bacteria is correlated with a cost for the host vector through the activation of an immune response and a developmental delay in the passage from L3 stage larvae to pupa (21–23). Moreover, overexpression of *evf* leads to *Drosophila* death (19, 21), suggesting that regulation of *evf* is essential not to kill the insect host before *Erwinia* can be transmitted to a new plant host.

By studying the tripartite interaction pathogen/plant host/insect vector our earlier work demonstrated that *evf* expression is triggered also during induction of the plant virulence factors responsible for maceration of plant tissue (21). This occurs due to co-regulation of *evf* and PCWDE expression by quorum sensing (21). This activation occurs in the absence of any insect cue, even though *evf* is necessary for the interaction with *Drosophila* but not in the plant infection. Coupling expression of virulence traits that cause degradation of plant tissues with the factor required for infecting the insect, could function as a mechanism to boost transmission. This would be akin to a predictive-like behavior (24–26), and could increase the probability of the bacteria surviving inside the insect attracted by the rotten plant tissue.

As virulence strategies often depend on functional associations between multiple genes, the commitment for expression of bacterial virulence can be costly, and thus is often regulated by multiple independent pathways (27, 28). In *Ecc15 evf* is regulated by quorum sensing via the global regulator RsmA and the transcriptional regulator Hor (20). Apart from quorum sensing no other pathways were known to regulate expression of *evf*. Therefore, we performed a genetic screen to identify novel transcriptional regulators of *evf*. This strategy allowed us to identify the sensor histidine kinase ArcB and a new TetR-like repressor, here named *lvtR* (**L**ow **V**irulence **T**ranscriptional **R**epressor), as novel regulators of *evf*. We show that these genes regulate *evf* and PCWDE through regulation of *hor* expression, independently of quorum sensing, and impact *Ecc15* pathogenesis in plants and *Drosophila*. We established a new infection assay where *Ecc15* causes lethality of *Drosophila* larvae, and we show that mutants in *arcB* and *lvtR* have the same survival rates as non-infected larvae. Moreover, we show that the ArcB role in regulating traits important for microbe-host interactions is conserved in other bacteria. Taken together, we reveal a central role of ArcB, LvtR and quorum sensing in the co-regulation of specific plant and insect factors and subsequent establishment of Plant-*Ecc15*-*Drosophila* interactions.

## Results

### Identification of *arcB* and *lvtR* as regulators of *evf* expression

To identify new regulators of virulence in *Erwinia*, we constructed a library of random Tn5::*kan* transposon (Lucigen) insertions in a wild-type (WT) *Ecc15* strain carrying an *evf* fluorescence reporter fusion (P*_evf_*::*gfp*). We screened 4800 mutants and selected those with lower P*_evf_*::*gfp* expression levels when compared to WT *Ecc15* at 6 hours of growth, corresponding to entry into stationary phase (Fig. S1). The full library was measured once, and the 10% mutants with lower or higher P*_evf_*::*gfp* expression in comparison the WT strain were rescreened. Following four repeated measurements 37 mutants showing low and 14 mutants showing high P*_evf_*::*gfp* expression levels in at least 3 out of the 4 measurements, were selected for identification of the transposon insertion site by whole genome sequencing (Table S1). To exclude mutants affected in bacterial growth, thus likely to be affected by pleiotropic effects in production of virulence, we measured the growth rate of these mutants in comparison to the WT strain. Likewise, to exclude mutants generally affected in transcription or biosynthesis potential, we calculated the ratio of P*_evf_*::*gfp* in relationship to the ratio of a constitutive *mCherry* fusion expressed in the same plasmid using the following formula: [(Mutant P*_evf_*::*gfp* level ÷ WT P*_evf_*::*gfp*) ÷ (Mutant P*_tet_*::*mCherry* ÷ WT P_tet_::*mCherry*)]. The ratio of the two fluorescences allowed us to exclude mutants with pleiotropic effects that, for instance, have lower levels of *evf* due to a general decrease in gene expression, as these will also show low amounts of *mCherry*. As cutoffs, we excluded mutants with effects in growth rate higher than 20% and ratios of P*_evf_*::*gfp* / P*_tet_*::*mCherry* higher than 90%. Using these criteria, and within the mutants with low *evf* expression, 16 mutants were selected (Table S2). As expected, these mutants include insertions in genes previously described as being involved in the regulation of *evf* (21), such as the gene encoding the quorum sensing signal synthase, ExpI, or the transcriptional regulator Hor (Table S2).

We have previously shown that ExpI regulates *evf* expression through the production of AHLs and, therefore, its mutant phenotype can be rescued in media supplemented with exogenous AHLs. Thus, we investigated if any of the 16 selected mutants could be rescued with exogenous AHLs. Exogenous supplementation of AHLs rescue P*_evf_*::*gfp* expression to WT levels only in the *expI* mutant (Table S2). Indicating that the new mutants do not reduce *evf* expression by affecting AHLs production.

One of the new regulators of *evf* expression that we identified is the gene *sapC*, involved in the transport of potassium. Interestingly, this gene was previously identified in a screen for regulators of PCWDE, where it was shown that active transport of potassium is important for the regulation of virulence (29). This result shows that screening for regulators of *evf* can identify PCWDE regulators as well. Therefore, to distinguish between specific regulators of *evf* and general regulators of virulence, we analyzed the expression levels of the PCWDE promoter *pelA* by measuring activity of a reporter fusion (P*_pelA_*::*gfp*), in all selected mutants. Using this approach, we observed that two mutants show altered expression of *evf* but have *pelA* expression levels similar to WT. These specific regulators of *evf* were *glpR*, the regulator of glycerol-3-phosphate metabolism, and the RNA chaperone *proQ*. All other mutants were affected in both *evf* and *pelA* expression, suggesting they are general regulators of virulence.

Overall, these results show that, by using this genetic screen approach, we were able to successfully identify novel regulators of *evf* expression. Out of the 16 mutants, two stand out which dramatically affect the expression of the *evf* reporter (85 and 75% reduction, respectively), this reduced *evf* expression is not complemented by AHLs, and their growth rate is not significantly affected (7 and 8%, respectively) (Table. S2). We selected these two mutants, carrying insertions in the genes *arcB* and 4038 (renamed herein *lvtR*, after **L**ow **V**irulence **T**ranscriptional **R**epressor), for further characterization.

### ArcB and LvtR are general regulators of virulence and affect the interaction of *Ecc15* with both plant and insect hosts

Following the results obtained with the transposon mutants, we constructed clean deletions of *arcB* and *lvtR* by allelic replacement in the *Ecc15* WT background. We confirmed that neither the deletion of *arcB* nor *lvtR* affect the growth of *Ecc15* (Fig. S1). Next, we measured the expression of both P*_evf_*::*gfp* and P*_pelA_*::*gfp* reporter fusions in WT, *arcB*, and *lvtR* mutants. We found that in the WT background both reporter fusions peak at 7 hours of growth (Fig. S2A-B). Therefore, to determine the impact of *arcB* and *lvtR* in the expression of virulence, we measured the expression levels of P*_evf_*::*gfp* and P*_pelA_*::*gfp* in each genotype at 7 hours of growth. We compared these results with those of an *expI* mutant, unable to produce AHLs, which we previously showed to have reduced *evf* and *pelA* expression. Consistent with the results obtained with the transposon mutants, both *arcB* and *lvtR* deletion mutants show lower levels of P*_evf_*::*gfp* expression compared to the WT (Fig. 1A, TukeyHSD test *p*<0.001, Fig. S2C-E). Both mutants also show lower levels of P*_pelA_*::*gfp* when compared to the WT (Fig. 1B, TukeyHSD test *p*<0.001, Fig. S2F-H), and as low as the *expI* mutant, suggesting that *arcB* and 4038 are general regulators of virulence in *Ecc15* possibly involved in both plant and insect infections.

**Fig. 1.**
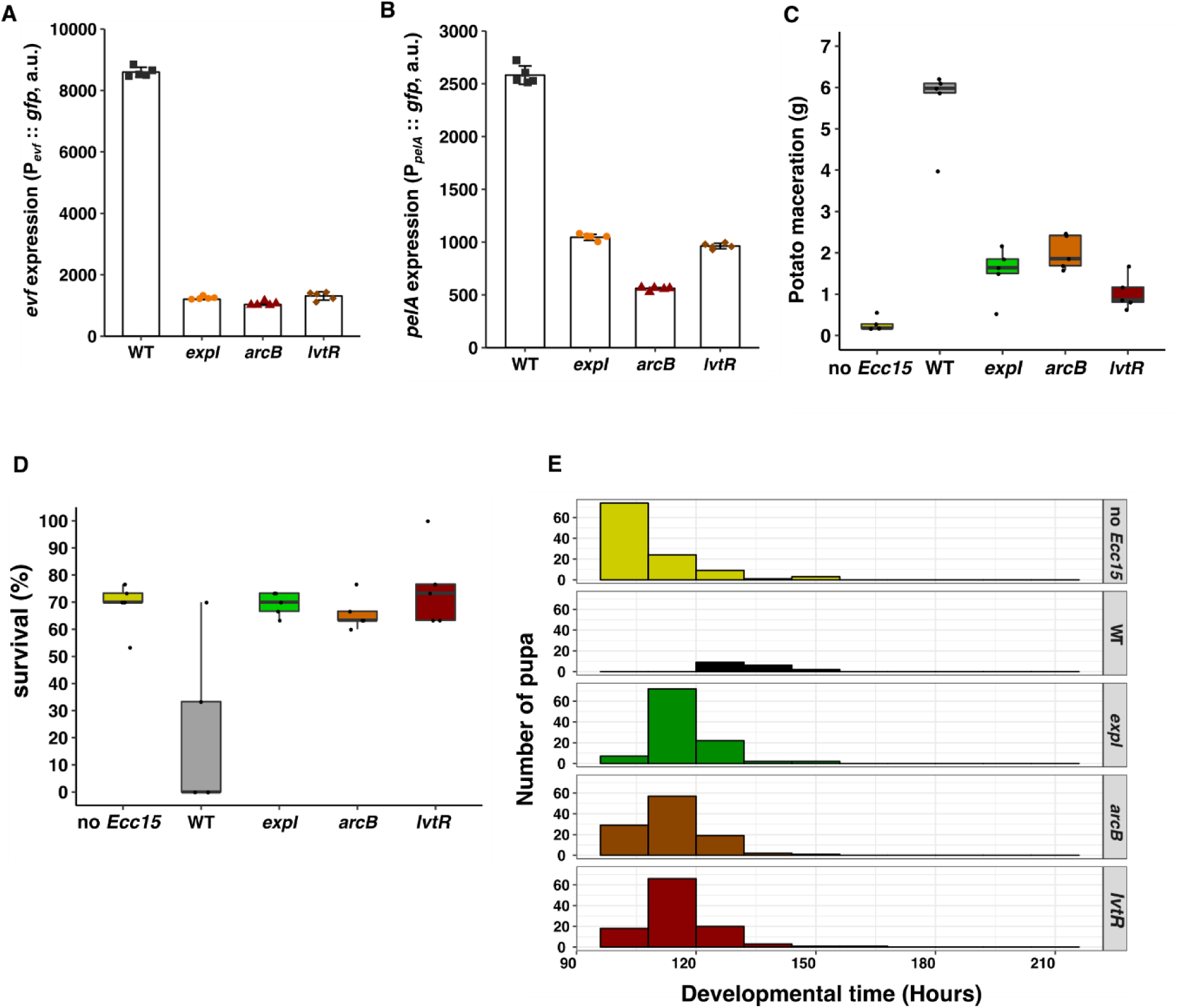
ArcB and LvtR regulate expression of virulence factors and are necessary for *Ecc15* infection phenotypes in plants and *Drosophila melanogaster*. **(A)** Pevf::*gfp* expression in WT *Ecc15*, *expI, arcB* and *lvtR* mutants at 7 hours of growth in LB + PGA 0.4% PGA + Spec. n=5 **(B)** PpelA::*gfp* expression in WT *Ecc15*, *expI, arcB* and *lvtR* mutants at 7 hours of growth in LB + PGA 0.4% + Spec. n=5 **(C)** Potato maceration quantification (grams) in potatoes infected with WT *Ecc15*, *arcB*, *lvtR* and mutants, 48 hours post-infection. n=5 **(D)** Survival measured as percentage of embryos that reach the pupa stage after exposure to WT *Ecc15, expI*, *arcB* and *lvtR* **(E)** *Drosophila* pupariation time after exposure of embryos to WT *Ecc15*, *expI*, *arcB* and *lvtR* compared with no *Ecc15*. Error bars represent standard deviation. For each panel a representative experiment from three independent experiments is shown (other two experiment are shown in Fig. S3). Statistical analysis with the data of all the three experiments is shown in Fig. S3.

To assess the impact of lacking either *arcB* or *lvtR* on the establishment of *Ecc15*-plant interactions, we measured the ability of these mutants to infect and macerate tissues of potato tubers. As previously described, the *expI* mutant shows lower levels of maceration when compared to the WT (Fig. 1C, TukeyHSD test p<0.001, Fig. S2I-K). The *arcB* and *lvtR* mutants also cause less maceration than the WT (TukeyHSD test *p*<0.001, FigS2K), and as low as the *expI* mutant (Fig. 1C, TukeyHSD test *p*=1, Fig. S2I-K). These results show that both *arcB* and *lvtR* are necessary for full pathogenicity of *Ecc15* towards the plant host.

*Ecc15* causes a developmental delay in *Drosophila melanogaster* L3 stage larvae that is dependent on *evf*, but is not known to affect the survival of these larvae (21, 22). Here, we established a new assay where we measured the impact of *Ecc15* in both survival and development time of *D. melanogaster* from early larvae to pupae. We added a bacterial suspension at a lower or higher density (1 and 100 OD, respectively) to *D. melanogaster* embryos and monitored their development till pupae. We found that a lower dose of *Ecc15* delayed development, but had survival rates similar to control (Fig. S3A-B). Interestingly, when a high dose of WT *Ecc15* was added there is high lethality and only 20% of the embryos reach the pupa stage (Fig. 1D, TukeyHSD test *p*<0.001, Fig. S2L-N). Moreover, we also observed that surviving larvae are 19 hours delayed in pupariation, on average, when compared to non-infected larvae (Fig. 1E, TukeyHSD test *p*<0.001, Fig. S2O-Q). Additionally, these phenotypes were observed with *Ecc15* but not with bacteria not carrying *evf*, such as *E. coli* or the *Ecc15* closely related species *Pectobacterium wasabiae strain Scc3193*, independently of the dose (Fig. S3A-B). Importantly, we found that larvae exposed to either *expI*, *arcB* or *lvtR* mutants have similar survival rate to non-infected larvae (Fig. 1D, TukeyHSD test *p*=1, Fig. S2L-N), and show no significantly developmental delay when compared to non-infected larvae (Fig. 1E, TukeyHSD test *p*=0.14, *p*=0.6 and *p*=0.11 respectively, Fig. S2O-K). Together, our results highlight the importance of *arcB* and *lvtR* as regulators of traits necessary for the establishment of *Ecc15* interactions with both the plant and the *Drosophila* vector, because *arcB* and *lvtR* mutants are avirulent in the plant assay and cause no developmental delay or *Ecc15* associated lethality in infected *Drosophila* larvae.

### ArcB and LvtR regulate production of the conserved regulator Hor independently of quorum sensing

As in *Ecc15* expression of *evf* is dependent on Hor (20, 21) we inquired if the lower levels of *evf* expression observed in the *arcB* and *lvtR* mutants were caused by downregulation of *hor* expression. We measured the levels of a *hor* by measuring fluorescence of a reporter fusion (P*_hor_*::*gfp*) in both *arcB* and *lvtR* mutants with respect to the WT. As shown before, the highest expression of the fusion in the WT is at 7 hours of growth (Fig. S4A), and P*_hor_*::*gfp* levels are lower in an *expI* mutant than in WT *Ecc15* (Fig. 2A, TukeyHSD test *p*<0.001, Fig. S4C-E). Moreover, we found that both *arcB* and *lvtR* mutants have P*_hor_*::*gfp* expression similar to an *expI* mutant and lower levels than WT *Ecc15* (Fig. 2A, TukeyHSD test *p*<0.001, Fig. S4C-E). These results suggest that the decreased *evf* expression observed in *arcB* and *lvtR* mutants are mediated by downregulation of *hor* expression.

**Fig. 2.**
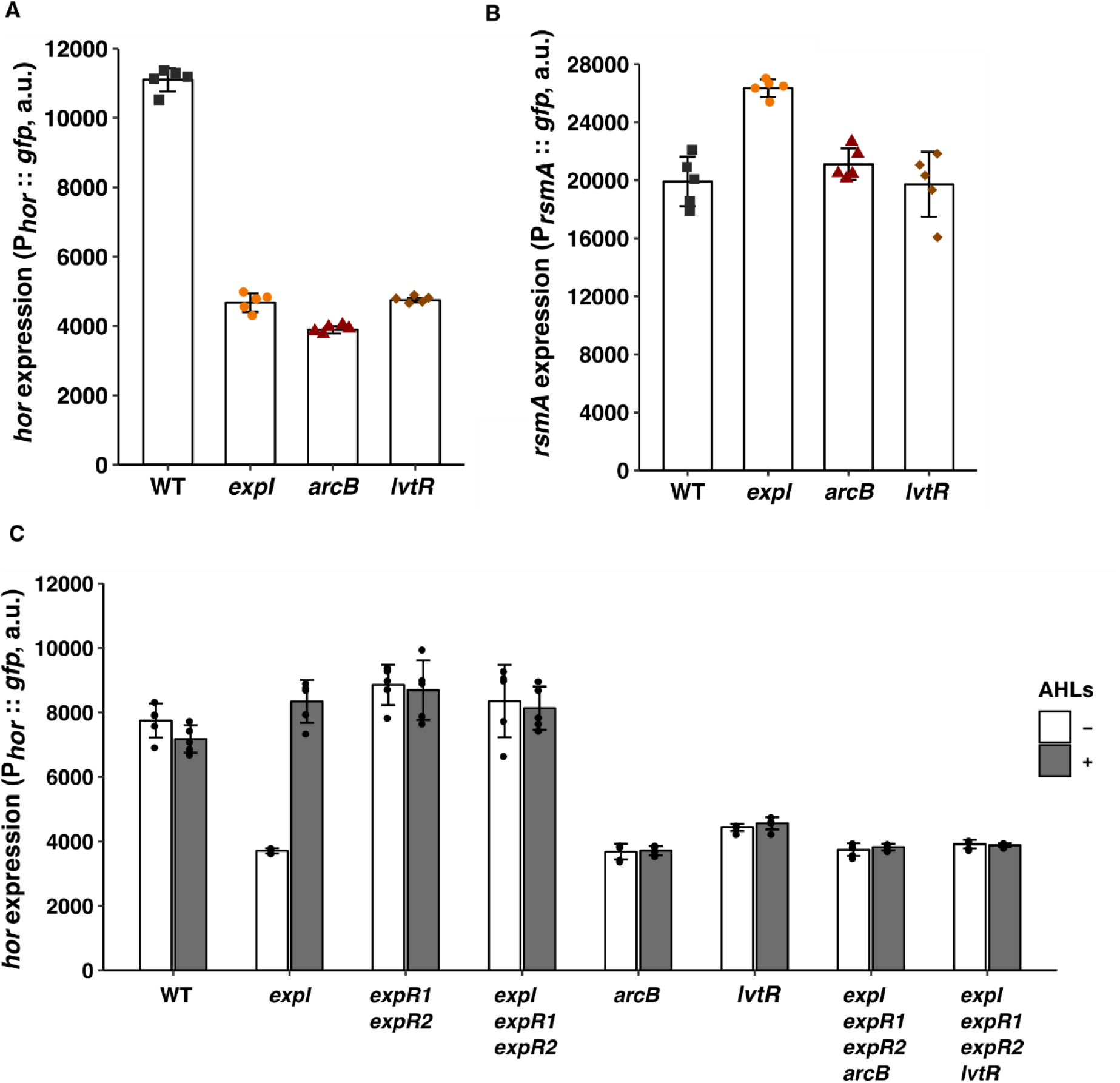
ArcB and LvtR regulate expression of *hor* independently of quorum sensing. **(A)** Phor::*gfp* expression in WT *Ecc15*, *expI, arcB* and *lvtR* mutants at 7 hours of growth in LB + PGA 0.4% + Spec. n=5 **(B)** PrsmA::*gfp* expression in WT *Ecc15*, *expI, arcB* and *lvtR* mutants at 3 hours of growth in LB + PGA 0.4% + Spec. n=5 **(C)** Phor::*gfp* expression without (white) or with (grey) addition of exogenous AHLs in WT *Ecc15*, *expI*, *expR1 expR2*, *expI expR1 expR2*, *arcB*, *lvtR*, *expI expR1 expR2 arcB* and *expI expR1 expR2 lvtR* mutants at 7 hours of growth in LB + PGA 0.4% + Spec. n=5. Complementation with AHLs was performed with a mixture of 1µM 3-oxo-C6-HSL and 3-oxo-C8-HSL.Error bars represent standard deviation. For each panel a representative experiment from three independent experiments is shown (other two experiment are shown in Fig. S4). Statistical analysis with the data of all the three experiments is shown in Fig. S4.

The transcription of Hor is regulated by the global negative regulator RsmA, which is in turn regulated by quorum sensing (30). So, we next asked if *arcB* and *lvtR* also regulate *rsmA* expression. To test this hypothesis, we measured the fluorescence of an *rsmA* reporter fusion (P*_rsmA_*::*gfp*) in both *arcB* and *lvtR* mutants. We found that expression of P*_rsmA_*::*gfp* is maximum at 3 hours of growth (Fig. S4B) and, we thus compared the levels of the fusion in the different mutants at that time point. As shown before, the *expI* mutant shows higher levels of P*_rsmA_*::*gfp* expression than the WT (Fig. 2B, TukeyHSD test *p*=0.005, Fig. S4F-H). However, both *arcB* and *lvtR* mutants show P*_rsmA_*::*gfp* levels similar to WT *Ecc15* (Fig. 2B, TukeyHSD test *p*=1, Fig. S4F-H), indicating that ArcB and LvtR regulate expression of virulence through regulation of *hor*, independently of RsmA. Since RsmA is strongly regulated by quorum sensing in this bacterium (12), these results also suggest that ArcB and LvtR regulate expression of virulence independently of quorum sensing. To further investigate this possibility, we measured *hor* expression in *arcB* and *lvtR* mutants introduced in an *expI expR1 expR2* background, which lacks both the synthase and the quorum sensing signal receptors, and is therefore blind to quorum sensing regulation. Both *arcB* and *lvtR* deletions in the *expI expR1 expR2* background still strongly affected the levels of P*_hor_*::*gfp* expression (Fig. 2C, TukeyHSD test *p*<0.001, Fig. S4I-K), to levels similar to *arcB* and *lvtR* single mutants (Fig. 2C, TukeyHSD test *p*=1, Fig. S4I-K). Importantly, we found that *expI* was the only mutant where expression of P*_hor_*::*gfp* expression responded to the exogenous supplementation of growth media with AHLs (Fig. 2C, TukeyHSD test *p*<0.001, Fig. S4I-K), whereas AHLs did not alter the effect of *arcB* and *lvtR* mutations on the expression of P*_hor_*::*gfp*. Taken together, our results show that ArcB and LvtR are required for the expression of *hor* and their effect is independent of the quorum sensing system. Furthermore, these results highlight a central role of Hor as an integrator of multiple signals.

### LvtR functions as TetR-like regulator controlling *hor* expression by repressing *lvhR*

Analysis of the aminoacid sequence of *lvtR* lead us to identify a sequence of 58 aminoacids correspondent to a HTH-like conserved N-terminal DNA binding domain associated with the TetR regulatory family of proteins (TFR). The majority of described TFRs, in the absence of their ligand, repress the gene coded immediately upstream of its own coding sequence (31, 32). To test if *lvtR* acts as a conventional TetR-like repressor, we constructed a GFP promoter fusion with the promoter region of the gene upstream of *lvtR*, i.e. the gene annotated as 4037, and measured the GFP levels of this fusion in a *lvtR* mutant. Like for the P*_evf_*::*gfp* fusion, the peak of expression for the P*_4037_*::*gfp* reporter fusion was at 7 hours of growth (Fig. S5A). We found that, indeed, *lvtR* downregulates 4037, as the *lvtR* mutant shows higher expression of the P*_4037_*::*gfp* reporter fusion when compared to the WT *Ecc15* (Fig. 3B, TukeyHSD test *p*<0.001, Fig. S5C-E). Next, we investigated if the LvtR-dependent regulation of *hor* could be mediated by 4037, by deleting it and measuring the impact of mutant in the expression of the *hor* reporter fusion. We found no significant difference in P*_hor_*::*gfp* expression levels in the single 4037 mutant when compared to those of the WT *Ecc15* (Fig. 3C, TukeyHSD test *p*=0.1, Fig. S5F-H). However, the double mutant *lvtR* 4037 shows higher expression levels of *hor* than the *lvtR* single mutant (TukeyHSD test *p*<0.001, Fig. S5H), and similar levels to those of the WT (Fig. 3C, TukeyHSD test *p*=0.99, Fig. S5F-H), indicating that the LvtR-dependent regulation of *hor* occurs via repression of 4037. We also checked *hor* expression in a deletion mutant of the gene 4039, located immediately downstream from LvtR. We saw no effect of 4039 in P*_hor_*::*gfp* expression either as a single or a double mutant with *lvtR* (Fig. 3C, TukeyHSD test *p*=1, Fig. S5F-H), indicating no role of this gene in the LvtR-dependent regulation of *hor*. Overall, these results suggest that LvtR and 4037 (but not 4039) form a pair that regulate *hor* expression, where LvtR is a repressor of 4037, which subsequently is a repressor of *hor* expression. Therefore, we herein rename 4037 as **L**ow **V**irulence **H**or **R**epressor (*lvhR*). We propose that *lvtR* is a type I one component transcriptional regulator of the TetR family, since LvtR represses the expression of *lvhR*, the gene located immediately upstream from *lvtR*, as it is common for the TetR family of regulators.

**Fig. 3.**
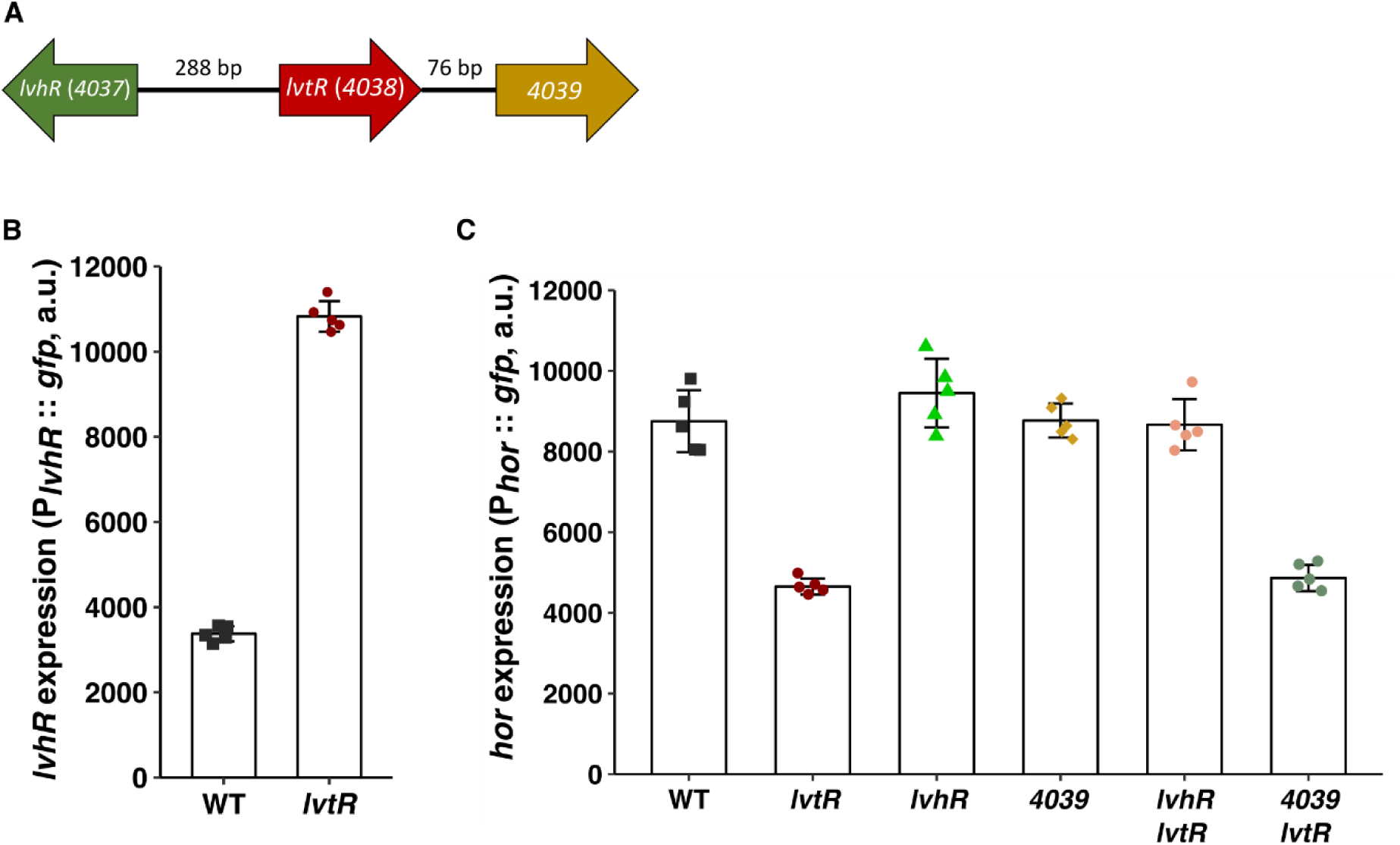
LvtR is a TetR-like response regulator that regulates *hor* through repression of *lvhR* expression. **(A)** Genomic organization of *lvtR* locus **(B)** P*lvhR*::*gfp* expression in WT *Ecc15* and *lvtR* mutants at 7 hours of growth in LB + PGA 0.4% + Spec. n=5 **(C)** Phor::*gfp* expression in WT *Ecc15*, *lvtR, lvhR*, *4039*, *lvhR lvtR* and *4039 lvtR* mutants at 7 hours of growth in LB + PGA 0.4% + Spec. n=5. Error bars represent standard deviation. For each panel a representative experiment from three independent experiments is shown (other two experiment are shown in Fig. S5). Statistical analysis with the data of all the three experiments is shown in Fig. S5.

### ArcB regulates the expression of *hor* independently of its cognate response regulator ArcA

In our genetic screen we identified ArcB as a regulator of virulence expression in *Ecc15*. Many proteobacteria carry homologues to ArcB. In *E. coli* ArcB/ArcA form a two component system that represses genes involved in aerobic respiration in response to decreasing levels of oxygen, which causes major changes in the redox state of the cell quinone pool (33). Coherently, ArcB/ArcA signalling was shown to be mostly active in microaerophilic conditions (34, 35). We asked if, in *Ecc15*, the regulation of *hor* expression by ArcB was dependent on oxygen levels. To test this, we measured the levels of P*_hor_*::*gfp* in bacteria growing with and without shaking, to generate higher and lower oxygen levels in the culture. We found no difference in the levels of P*_hor_*::*gfp* in the WT *Ecc15* nor in the *arcB* mutant grown under these two conditions (Fig. 4A, TukeyHSD test *p*=0.5, and *p*=0.8, Fig. S6A-C). Therefore, we found no evidence that ArcB*-*mediated regulation of *hor* is dependent on oxygen levels in *Ecc15*. In *E. coli*, activation of ArcB is favored by a change in the redox state of the pool of ubiquinone and menaquinone from majorly oxidized to majorly reduced (36, 37). To further ratify that the decrease of *hor* expression observed in the *arcB* mutant is independent of oxygen and redox state regulation, we constructed two mutants affected in the production of quinones. The *ubiC* mutant is unable to convert chorismate to 4-hydroxybenzoate and pyruvate, an essential step for the production of ubiquinone, and the *menF* mutant is affected in the conversation of chorismate to isochorismate essential for the production of menaquinone. We analyzed the levels of *hor* expression in these two mutants in higher and lower oxygen levels (Fig. S6D). While P*_hor_*::*gfp* in the *arcB* mutant is lower than the WT, we observed no difference in P*_hor_*::*gfp* expression between the WT and the *ubiC* or *menF* mutants growing in either higher or lower levels of oxygen (Fig. S6D). This reinforces the evidence that, in *Ecc15*, ArcB is active in the presence of oxygen, and regulates expression of *hor* independently of oxygen levels.

**Fig. 4.**
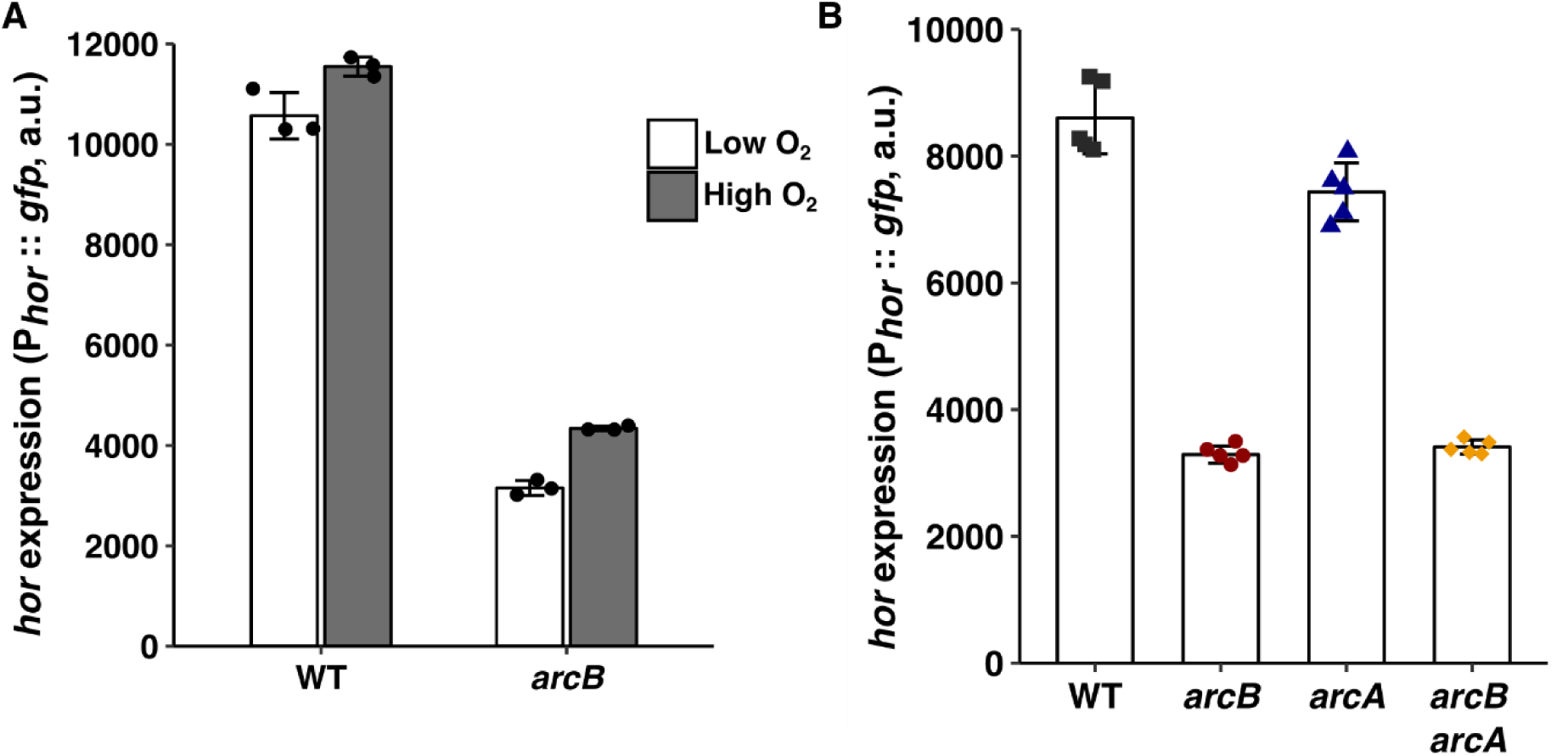
ArcB regulates expression of *hor* independently of oxygen levels and the cognate response regulator ArcA. **(A)** Phor::*gfp* expression without (white) and with (grey) shaking to promote a decrease in oxygen availability in WT *Ecc15* and *arcB* mutants at 7 hours of growth in LB + PGA 0.4% + Spec. n=3 **(B)** Phor::*gfp* expression in WT *Ecc15*, *arcB, arcA* and *arcB arcA* mutants at 7 hours of growth in LB + PGA 0.4% + Spec. n=5. Error bars represent standard deviation For each panel a representative experiment from three independent experiments is shown (other two experiment are shown in Fig. S6). Statistical analysis with the data of all the three experiments is shown in Fig. S6.

The ArcB/ArcA two component system was previously shown to promote bacterial resistance to reactive oxygen species (38, 39). Therefore, to test if the ArcB/ArcA two component system of *Ecc15* shares functionality with those of other bacterial species, we also generated a mutant in *arcA*, and tested the role of ArcB/ArcA in resistance of *Ecc15* to hydrogen peroxide. We found that single *arcB*, single *arcA*, and *arcB arcA* double mutants, are more susceptible to H_2_O_2_ than the WT *Ecc15* (Fig. S6I), suggesting that the ArcB/ArcA two component system of *Ecc15* shares functionality with that of *E. coli.* Next, we tested if the decrease of P*_hor_*::*gfp* expression observed in the *arcB* mutant is dependent on its cognate response regulator (RR) ArcA. To test this, we measured the levels of *hor* expression in both an *arcA* and *arcB arcA* double mutant. We found no difference in P*_hor_*::*gfp* expression levels in an *arcA* mutant when compared to the WT *Ecc15* (Fig. 4B, TukeyHSD test *p*=0.1, Fig. S6F-H). Consistently with these results, we found no significant differences in P*_hor_*::*gfp* expression levels in the double *arcB arcA* mutant when compared to the *arcB* single mutant (Fig. 4B, TukeyHSD test *p*=1, Fig. S6F-H). Therefore, we concluded that, although ArcB requires its cognate RR ArcA for the regulation of resistance to H_2_O_2_, it does not require it for the regulation of *hor*.

### ArcB is a conserved regulator of multiple traits involved in host-microbe interaction

The ArcB/ArcA two component system is conserved among different bacterial species, many of them known to be involved in interactions with Eukaryotic hosts (38–40), but the potential role of ArcB in regulating traits required for these interactions has been poorly investigated. Given our finding that ArcB regulates virulence in *Erwinia*, we asked if it can also be important for the regulation of traits involved in establishing host-microbe interactions in other bacterial species. To test this, we constructed *arcB* deletion mutants in *Salmonella enterica* serovar Thyphimurium and *Vibrio harveyi*, and measured the impact of this mutation on the formation of biofilms and bioluminescence, respectively. Biofilm formation in *Salmonella* is characterized by the production of curli fibers, which gives a rough morphology to colonies when grown under oxygen tension, nutritional stress and osmotic pressure (41). Production of these fibers is regulated by the transcription factor CsgD, and mutants lacking this gene are characterized by showing a smooth colony morphology. As previously described, we observed a rough colony morphology in the WT, whereas colonies of the *csgD* mutant are smooth (Fig. 5A, Fig. S7A). Importantly, we found that an *arcB* mutant shows a smooth colony morphology similar to the *csgD* mutant, and very distinct from the WT strain (Fig. 5A, Fig. S7A). Next, to measure the capacity for biofilm formation, we quantified the amount of total biomass produced by WT and *arcB Ecc15*, by using a crystal violet staining assay. As expected, we found that a *csgD* mutant produces less matrix than the WT strain (Fig. 5B, TukeyHSD test *p*<0.004, Fig. S7B-D). Moreover, we found that an *arcB* mutant produces less matrix than both the *csgD* mutant and the WT strain (Fig. 5B, TukeyHSD test *p*<0.001, Fig. S7B-D). Next, we tested if deletion of *arcB* also affects production of bioluminescence in *Vibrio harveyi*, a trait essential for its establishment in the light organ of the Hawaiian bobtail squid *Euprymna scolopes* (42). Bioluminescence in *Vibrio harveyi* is controlled by quorum sensing, namely by the production of the AI-2 signal carried by the *luxS* gene (43, 44). We found that an *arcB* mutant shows lower levels of bioluminescence than the WT (Fig 5C, TukeyHSD test *p*<0.001, Fig. S7E-G), and as low as a *luxS* mutant (Fig 5C, TukeyHSD test *p*<0.001, Fig. S7E-G). Our results show that, beyond regulating virulence in *Ecc15*, ArcB also plays a major role in the regulation of biofilm formation in *Salmonella* and of bioluminescence in *Vibrio*, revealing that this histidine kinase is a conserved regulator of traits associated with the establishment of host-microbe interactions.

**Fig. 5.**
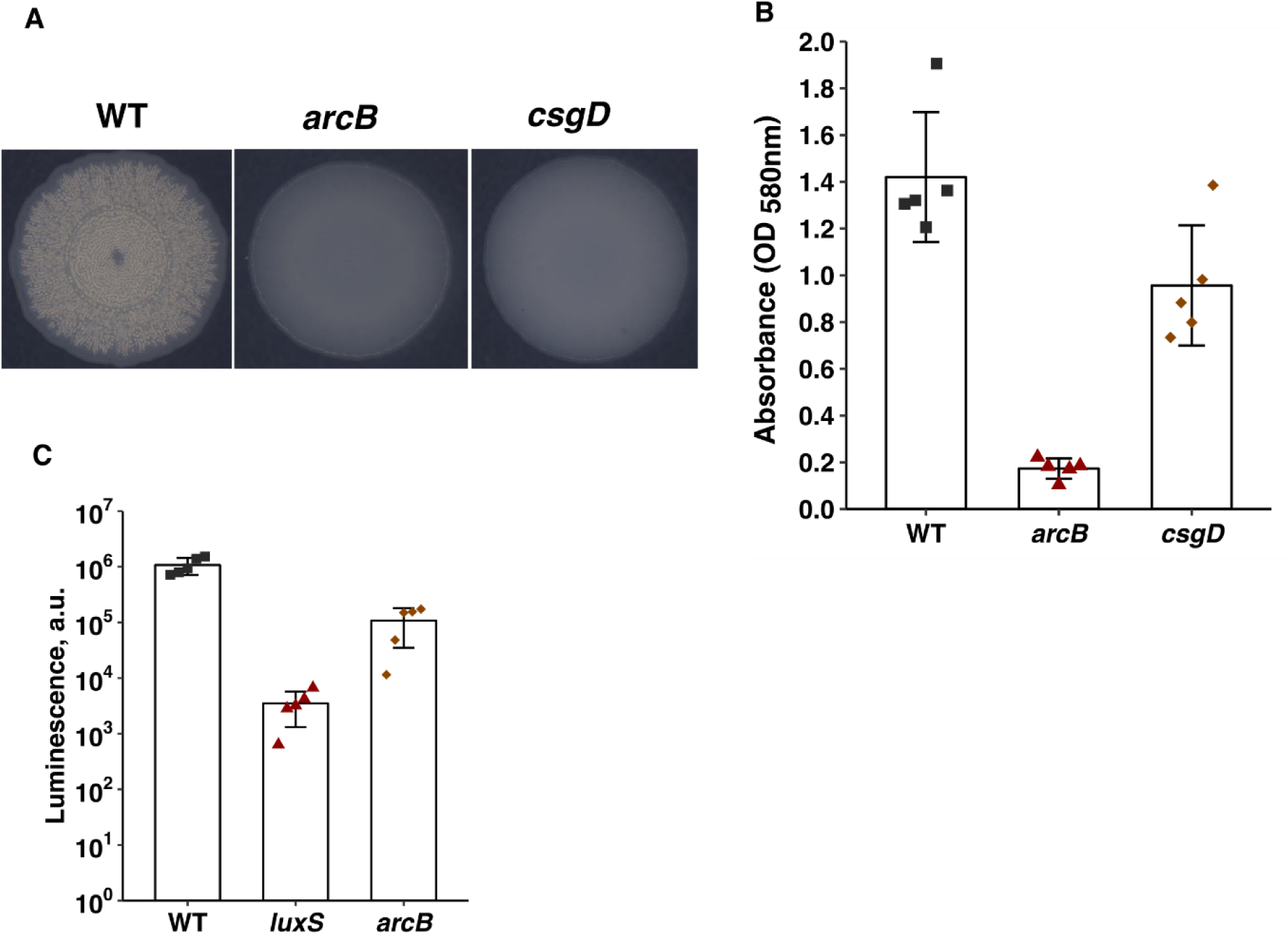
ArcB regulates formation of biofilms and luminescence in *S.* Typhimurium and *V. harveyi*. **(A)** Colony morphology of WT, *arcB*, and *csgD* mutants of *S.* Thyphimurium grown in LB without salt at 28°C **(B)** Quantification of biofilm formation in WT, *arcB*, and *csgD* mutants of *Salmonella* grown in LB without salt at 28°C. n=5. Error bars represent standard deviation of the mean. **(C)** Quantification of luminescence in WT, *luxS* or *arcB* mutants of *V. harveyi* grown in AB at 30°C. n=5. Error bars represent standard deviation. For each panel a representative experiment from three independent experiments is shown (other two experiment are shown in Fig. S7). Statistical analysis with the data of all the three experiments is shown in Fig. S7.

## Discussion

In *Ecc15*, expression of PCWDE is tightly controlled by the dynamic integration of information on cell density via the quorum sensing system and on environmental cues, namely by detection of plant metabolites, such as polygalacturonate, a component of the plant cell wall (14, 45, 46). Expression of *evf* is also regulated by quorum sensing, but it was not known if additional regulatory mechanisms converge to control the production of this virulence factor. Using a forward genetic screen, we identified ArcB and a TetR-like repressor, 4038, herein renamed LvtR, as novel regulators of *evf*. This suggests that, like PCWDE, expression of *evf* also responds to the integration of multiple signaling cues. Additionally, we found that these regulators also control expression of *pelA*, one of the PCWDE, revealing extra layers of regulation for the production of PCWDE as well. Accordingly, *arcB* and *lvtR* mutants are unable to macerate the tissues of plant tubers or to induce lethality in *Drosophila melanogaster* larvae, revealing the relevant role of these genes in the establishment of *Ecc15*-host-vector interactions.

We showed that ArcB and LvtR control PCWDE and *evf* expression via regulation of *hor* expression. Hor is a conserved transcription regulator essential for activating expression of *evf* and PCWDE (20, 21) and regulated by quorum sensing via the *expI*/*expR* system (21, 47, 48). Briefly, at low cell densities, the ExpR1 and ExpR2 quorum sensing receptors function as transcription activators, promoting the expression of the global negative regulator RsmA which consequently represses expression of PCWDE by inhibiting Hor expression (12). At high cell density, the ExpR1 and ExpR2 quorum sensing receptors bind to homoserine lactones, losing their ability to bind DNA, which leads to downregulation of *rsmA* expression, increase expression of Hor, activation of Evf and PCWDE (30, 49–52, and Fig. 6). RsmA is a conserved post transcriptional regulator that has been the focus of many studies on regulation of expression of virulence and secondary metabolism in different bacterial species (53, 54). In *Erwinia*, RsmA has been proposed to function as a signaling integration hub that receives input from the quorum sensing system, and from the environment via KdgR, a RR thought to respond to the presence of polygalacturonate (14, 15, 46). Therefore, we were interested in understanding if ArcB and LvtR regulate *hor* expression independently of RsmA. We showed that neither *arcB* nor *lvtR* mutants influence *rsmA* expression, suggesting that ArcB and LvtR-dependent regulation of *hor* is not mediated by RsmA. Since *rsmA* expression is regulated by quorum sensing (Fig. 3B and (12)), and it is thought to be required to mediate the quorum sensing response of *hor*, these results indicate that regulation of *hor* by ArcB and LvtR is independent of quorum sensing. Therefore, to test this we analyzed the effect of removing *arcB* and *lvtR* in a mutant background blind to quorum sensing, the *expI expR1 expR2* triple mutant. We found that in both the *arcB expI expR1 expR2* and *lvtR expI expR1 expR2* quadruple mutants the levels of *hor* expression are lower than the *expI expR1 expR2* triple mutant, and as low as the *arcB* or *lvtR* single mutants. Together, these data show that both ArcB and LvtR regulate the expression of *hor* independently of quorum sensing, suggesting that, in *Ecc15*, Hor functions as a signaling hub, integrating signals coming both from the *expI/expR* quorum sensing system and environmental sensing via LvtR and ArcB.

**Fig. 6.**
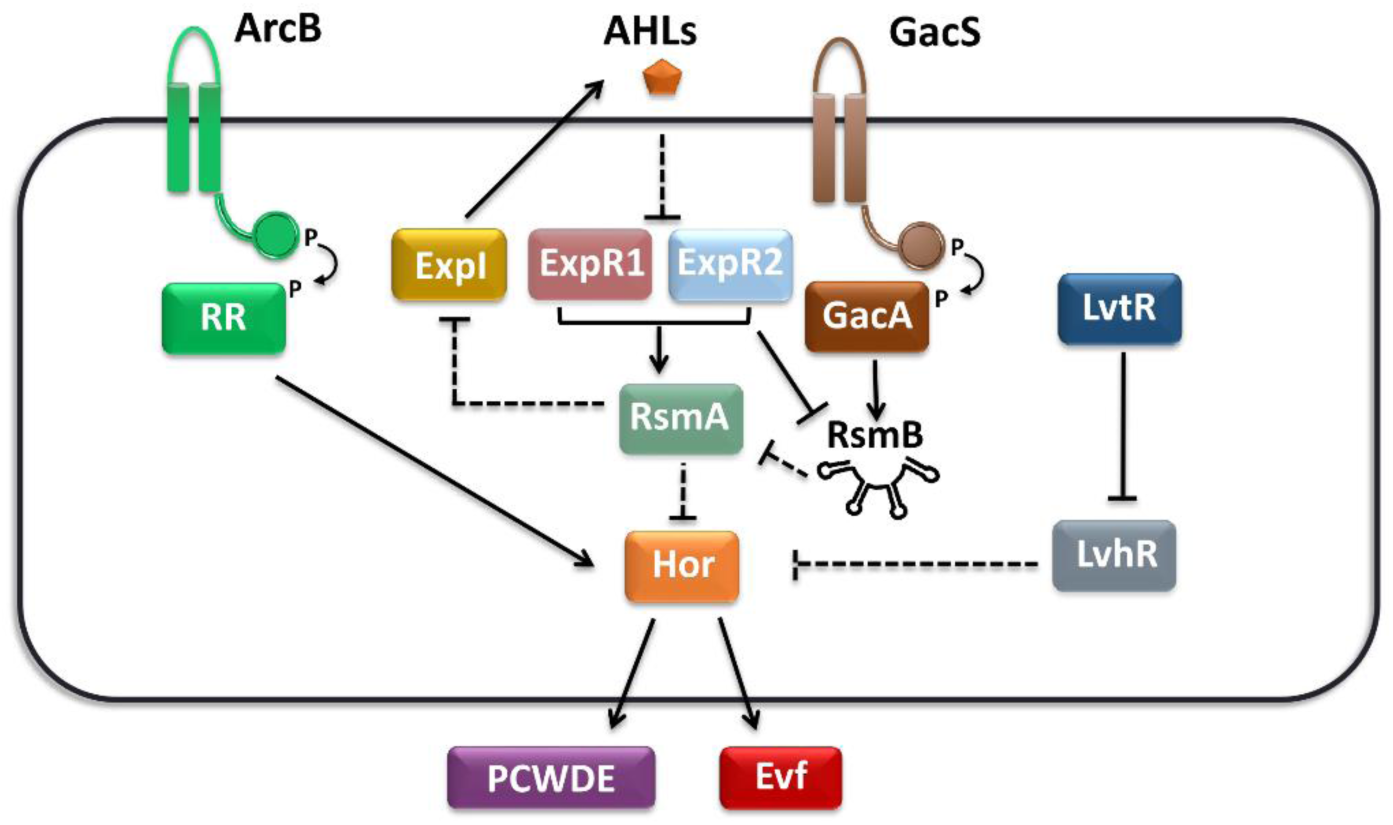
Integration of quorum sensing, ArcB and LvtR signalling for the control of *hor* transcription and consequently PCWDE and *evf* in *Ecc15*. *Ecc15* the synthase EpxI produces AHLs. As cell density increases, AHLs accumulate and when the concentration threshold is reached these signal molecules bind to ExpR1 and ExpR2 receptors inhibiting their ability to bind DNA. As ExpR1 and ExpR2 are required to induce *rsmA* transcription, expression of RsmA decreases. The GacS/A two component system is also active at high cell density and promotes transcription of *rsmB*, a noncoding RNA that binds the remaining available RsmA inhibiting it. Low levels of RsmA will lead to derepression of *hor* expression, the consequence increase in Hor levels leads to expression of virulence. We show here that Hor is also regulated by ArcB and the TetR family regulator *lvtR*. Taken together our data suggests that Hor functions as a signaling hub integrating the input of the quorum sensing system via RsmA, of ArcB via an unknown RR and of LvtR via LvhR. Arrows indicate activation, while intersecting lines indicate repression. Solid lines indicate transcriptional regulation, while dashed lines indicate post-transcriptional and post-translational mechanisms.

We also identified LvtR as a transcription regulator of both *evf* and *pelA* in *Ecc15.* This regulator is present in different *Erwinia* species, has an N-terminal DNA binding motif conserved among the tetracycline family of repressor genes (TFR), and it acts as a typical one-component RR. The one-component RR are a class of signal transduction transcriptional regulators typically involved in environmental sensing and are widely distributed among bacterial species (55, 56). These proteins are characterized by containing both a sensory domain, and either a C-terminal, or an N-terminal DNA binding domain. These regulators can bind to a variety of different ligands, and function either as activators or repressors of gene expression (31). TetR, involved in repression of the *tet* operon, which confers resistance to the antibiotic tetracycline, it is the best described member of one-component systems giving its name to the TFR family of proteins (57). TetR-like regulators are expected to bind specific cognate ligands, however most of the these ligands are unknown (32). The TFRs genomic architecture is usually conserved and, in most of the cases, they act on the gene coded upstream of their own coding sequence. We showed that, in *Ecc15*, LvtR acts as a typical TFR because it represses expression of the upstream gene, here *lvhR*, which in turn represses *hor* expression. LvhR is a hypothetical protein that seems to be conserved among different bacterial species. Interestingly, LvhR possesses a protein folding and a conserved ATP-binding domain similar to that of the plant gene TM-1, which confers resistance to infections by the tomato mosaic virus (ToMV). TM-1 inhibits viral replication by binding to viral RNA proteins involved in translation (58). This binding was shown to be dependent on the N-terminal domain of TM-1 and on the presence of ATP (59). This suggests that LvhR may regulate expression of *hor* by inhibiting a regulator through protein-protein interactions and not by directly regulating its expression via DNA binding. Considering that LvhR functions as a repressor of *hor* expression, our results indicate that the LvtR ligand is not present in our experimental conditions, thus keeping *lvhR* repressed during cell growth and allowing for the expression of *hor*. A *lvtR* mutant is unable to produce both PCWDE and *evf*, rendering the bacterium avirulent. Because this gene seems to be conserved among *Erwinia* species, identification of a LvtR ligand would be of great biotechnological interest as an alternative target for the development of efficient therapies to control *Erwinia* pathogenicity and, potentially, that of other bacterial pests. Further studies are necessary to understand both the molecular mechanism of *hor* repression by *lvhR*, as well as the nature of the LvtR ligand.

While one-component systems are typically limited to sensing compounds present in the cytosol, two-component systems (TCS) offer the benefit of responding to external stimuli at the level of the cell membrane (55). These systems are composed by a membrane anchored histidine sensor kinase (HK), like ArcB, and a respective RR. Upon sensing of the external stimulus, the HK autophosphorylates and subsequently phosphorylates the RR, which then will be active to induce or repress gene expression (55, 60–62). In our genetic screen we found that ArcB, the HK component of the ArcB/ArcA TCS, is necessary for expression of virulence in *Erwinia*. ArcB/ArcA belongs to a group of TCS denominated phosphorelays. In this type of TCS, the phosphoryl group is transferred multiple times across different domains of the HK before it reaches the RR (55). The ArcB/ArcA TCS is widely conserved and is very well studied in *E. coli*, where it is known to repress genes involved in aerobic respiration in response to decreasing concentrations of oxygen (63). It was previously shown in *E. coli*, that in response to a decrease in oxygen levels, there is a switch in the redox state of the quinone pool from majorly oxidized to the reduced form. This switch triggers autophosphosrylation of ArcB, which consequently phosphorylates ArcA, leading to repression of genes involved in aerobic respiration (36, 37, 64). However, in *Ecc15* we found no significant differences in the levels of *hor* expression in cells growing under higher or lower oxygen concentrations, suggesting that, in our experimental conditions, ArcB mediated regulation of *hor* is independent of oxygen. Moreover, although we showed that similarly to *E. coli* both the *arcB* and *arcA* are required to confer resistance to H_2_O_2_ in *Ecc15*, and thus providing evidence that this TCS also has a canonical role in this *Erwinia* strain, we also found that the regulation of *hor* by ArcB is not mediated by the cognate RR ArcA. As ArcA does not seem to play a role in the regulatory processes studied here, one possibility is that, in our experimental conditions ArcB affects phosphorylation of a non-cognate RR. Another possibility is the participation of ArcB in a multikinase network. These are groups of interconnected HK that directly affect the degree of phosphorylation of each other (61). For instance, RetS, a HK of *Pseudomonas aeruginosa*, directly affects the degree of phosphorylation of GacS. It was shown in this bacterium that RetS directly interacts with GacS and can promote its dephosphorylation, indirectly affecting the degree of phosphorylation of the RR GacA and consequently gene expression (8, 65). Even in *E. coli* there is accumulating evidence that ArcB can also phosphorylate non-cognate RR regulators. For instance, in *E. coli*, it was observed that while entering into stationary phase, even in a high-oxygen environment, ArcB can promote phosphorylation of both ArcA and the non-cognate RR RssB (66). It was hypothesized that the degree of specificity of ArcB-mediated phosphorylation depends to a certain extent on both oxygen and energy supply allowing for both a tight control over transcription of *rpoS*, which encodes σ^S^, the sigma factor controlling transition into stationary phase, and also its proteolysis. Also in *Erwinia amylora*, ArcB is up-regulated in response to copper toxic shock, and affects the ability of the bacterium to survive both *in vitro* and *in planta*, suggesting that cooper may be one of the signals that affect ArcB phosphorylation state (67). It is possible that during the infection of the plant, as bacterial cell density increases and both nutrients, metals and oxygen levels are depleted, ArcB triggers phosphorylation of a non-cognate RR in order to further activate expression of *evf*. Nevertheless, the full physiological implications of this regulation need further investigation.

The ArcB/ArcA TCS is conserved among different bacterial species and it has been shown to regulate expression of multiple traits (38–40, 68, 69). Particularly, it has been shown to regulate resistance of *E. coli*, *Haemophilus influenza* and *Salmonella enterica* to reactive oxygen and nitrogen species (39, 70, 71), a common antimicrobial feature of the immune response. We showed here that an *arcB* mutant of *Ecc15* is unable to cause tissue maceration of the plant host, as well as to cause a developmental delay in *Drosophila melanogaster*. Due to the large degree of conservation of ArcB/ArcA in both environmental and host-associated bacteria, we decided to test if ArcB has a conserved role in the regulation of biofilm formation in *Salmonella enterica* serovar Thyphimurium. Mammalian infections by *Salmonella* are often associated with consumption of contaminated food, including that of plant origin (72). Interestingly, biofilm formation is a key component of *Salmonella* survival strategy in both plants and in the gut of animals (73–76). The biofilm matrix is composed by different factors, including curli fimbriae, which are specifically produced by *Salmonella* and confer a rough morphology to colonies (77–79). Importantly, production of these fibers and, consequently biofilms, is controlled by the transcription factor CsgD in response to different environmental stimuli, such as oxygen tension, nutrient depletion and osmotic stress (41, 80). The regulation of *csgD* is highly complex, involving multiple regulatory proteins such as OmpR, IHF, H-NS, RpoS and CpxR (81). Interestingly, transcription of *csgD* during aerobiosis is promoted by the binding of phosphorylated OmpR to the promoter region of *csgD*, although phosphorylation is not dependent on the cognate HK EnvZ. We found that an *arcB* mutant of *Salmonella* shows similar phenoytes as a *csgD* mutant, such as smooth colony morphology and impaired production of cell matrix, essential for the formation of biofilms. One possibility is that activation of *csgD* is mediated by ArcB signalling via phosphorylation of OmpR. While this hypothesis still needs to be investigated, our results highlight the role of ArcB in the formation of biofilms in *Salmonella*. Moreover, the ArcB/ArcA TCS has been shown to regulate production of bioluminescence in *Vibrio fischeri*, a trait that affects colonization of the *Euprymna scolopes* light organ by this bacterium (42, 69, 82). Previously, it was shown that a mutant for *arcA* mutant produces more light, suggesting that ArcB/ArcA represses production of light (69). While the work done before was mostly focused on the RR ArcA, here we found that an *arcB* mutant of *Vibrio harveyi* produces less light than the wild type. This apparently contradictory result reinforces the notion of ArcB being capable of phosphorylating non-cognate RR. In that scenario, under certain environmental conditions (i.e. higher levels of oxygen) ArcB could regulate the phosphorylation state of a non-cognate RR, which could initiate a regulatory cascade leading to regulate production of light. Our data clearly indicates that ArcB has an active role in the establishment of several host-microbe interactions, and, in particular, in the regulation of virulence and establishment of *Ecc15*-host-vector interactions.

Our results show that *Ecc15* integrates environmental, cell-to-cell, and physiological cues to co-regulate expression of PCWDE and *evf*. Besides quorum sensing, we found that ArcB and LvtR regulate expression of virulence, through Hor. These proteins are putatively responding to environmental signals independently of quorum sensing thus making Hor an integrator of quorum sensing and other environmental signals in the regulation of virulence and colonization (Fig. 6). We hypothesize that ArcB is responding to the metabolic state of the cell, while LvtR binds to an unknown environmental cue not present in our experimental conditions. While *evf* promotes the interaction with the insect vector, in excess can be lethal for it, making regulation of Evf production an essential aspect of *Erwinia* lifestyle. Integration of diverse signals thus ensures that the expression of relevant genes is balanced and well timed. Moreover, signal integration allows co-expression of factors necessary in the plant stage of infection and those necessary in the vector stage, engaging in a predictive-like behavior likely advantageous for vector-borne pathogens, by favoring the interaction with the vector and, ultimately, spreading. Therefore, our work reinforces the idea that integration of multiple cues allows to combine fast response, fine-tune titration of gene expression, and maximization of resources to facilitate the interaction between microbes and multiple hosts.

## Material and Methods

### Bacterial strains, plasmids, and culture conditions

The strains and plasmids used in this study are listed in Table S3 of the supplementary material. All *Erwinia* strains used are derived from wild-type (WT) *Ecc15* strain. *Ecc15* and mutants were grown at 30°C with aeration in Luria-Bertani medium (LB). When specified, medium was supplemented with 0.4% polygalacturonic acid (PGA; Sigma P3850), to induce the expression of PCWDEs, or strains were grown without aeration to reduce oxygen availability. *E. coli* DH5α was used for cloning procedures, and *E.coli S17 λpir* to perform conjugation. Both were grown at 37°C with aeration in LB, unless specified. When required, antibiotics were used at the following concentrations (mg liter^−1^): ampicillin (Amp), 100; kanamycin (Kan), 50; spectinomycin (Spec), 50; chloramphenicol (Cm), 25, 10, 5; gentamycin (Gent), 15; Polymixin B (PB), 50. To assess bacterial growth, optical density at 600 nm (OD_600_) was determined in a Thermo Spectronic Helios delta spectrophotometer. Electro competent cells of both *Erwinia* and *Salmonella* were prepared by growing cells until OD_600_ ≈ 0,6 in LB supplemented with spec 50 or amp 100, respectively, and arabinose at 1 mM concentration to induce λ-Red recombineering system. Cells were then gently washed 3 times with glycerol 10% and pelleted by centrifugation for 20 min at 4000 rpm, in a previously cooled to 4°C centrifuge. After washes, cells were resuspended in 200 µl of 10% glycerol and kept in ice until further use. For conjugation, *Vibrio* was grown in LM at 30°C overnight, and *E.coli S17 λpir* at 37°C, both with aeration. 1ml of each culture was centrifuged and resuspended in 1ml of new media. 7 µl of each strain were then mixed, spotted in LB, and incubated overnight at 30°C.

### Genetic and molecular techniques

All primer sequences used in this study are listed in Table S4 in supplemental material. *Ecc15* deletion mutants listed in Table S2 were constructed by chromosomal gene replacement with an antibiotic marker using the λ-Red recombineering system (83). Plasmid pLIPS, able to replicate in *Ecc15*, and carrying the arabinose-inducible λ-Red recombineering system was used (9, 29). Briefly, the DNA region of the gene to be deleted, including approximately 500 bp upstream and downstream from the gene, was amplified by PCR and cloned into pUC18 (84) using restriction enzymes. These constructs, containing the target gene and its flanking regions, were divergently amplified by PCR, to introduce an *Xho*I restriction site in the 5′ and 3′ regions and to remove the native coding sequence of the target gene. The kanamycin cassette from pkD4 was amplified with primers also containing the *Xho*I restriction site. The fragment containing the kanamycin cassette was then digested with *Xho*I and was introduced into the *Xho*I-digested PCR fragment carrying the flanking regions of the target gene. The final construct, containing the kanamycin cassette flanked by the upstream and downstream regions of the target gene was then amplified by PCR, and approximately 2 micrograms of DNA fragment were electroporated into the parental strain (FDV31) expressing the λ-Red recombinase system from pLIPS, to favor recombination (9).

To construct the plasmid carrying the promoter *evf* fused to GFP (pFDV54), a fragment of 503 bp containing the *evf* promoter was amplified from WT *Ecc15* DNA with the primers P1194 and P1195. This fragment was then digested with *Hin*dIII and *Sph*I and ligated to pUC18. GFP was amplified from the pCMW1 (85) vector using primer P0576 and P0665. Both the GFP and pUC18-P_evf_ were digested with *Sph*I and *Bam*HI, ligated and 2 µl of the ligation reaction were used to transform Dh5α (pFDV54). The same procedure was used for the P*_hor_*::*gfp* fusion using primers P1351 and P1352 for promoter amplification (493 bp) and primers P1353 and P1354 for GFP amplification. Digestions were made with enzymes *Hin*dIII/*Pst*I and *Pst*I/*Xba*I (pFDV84). For P*_pelA_*primers P1941 and P1942 were used for promoter amplification (300 bp) and GFP was amplified using P1333 and P1334. Digestions were made using *Hind*III/*Xba*I and *Xba*I/*Sac*I. pOM1-mCherry was constructed by digesting pOM1 with *Xmn*I and ligating a fragment of 825 bp amplified with primers P1789 and P1790 from genomic DNA of the strain RB290 containing the constitutive mCherry fusion.

To construct the *arcB* mutant in *Salmonella* a kanamycin cassette flanked by a homologous region of 50 bp upstream and downstream of the *arcB* open reading frame was generated from pKD4 by PCR using primers P2070 and P2071. Approximately 2 micrograms of PCR amplified DNA were electroporated into the parental strain expressing the λ-Red recombineering system from pKD46. The *arcB* mutant in *Vibrio* was constructed using a modified version of the protocol from Ushijima et.al. (85). The DNA region of *arcB*, including approximately 500 bp upstream and downstream from the gene, was amplified by PCR with primers P2266 and P2267 from BB120 genomic DNA and cloned into pOM1 using *EcoRI/PstI* restriction enzymes. These constructs, containing *arcB* and its flanking regions, were divergently amplified by PCR with primers P2196 and P2197, to introduce a *XhoI* restriction site in the 5′ and 3′ regions and to remove the native coding sequence of *arcB* gene. The gentamicin cassette was amplified from strain RB980 with primers P1617 and P1618 also containing the *XhoI* restriction site. The fragment containing the gentamicin cassette was then digested with *XhoI* and was introduced into the *XhoI*-digested PCR fragment carrying the flanking regions of *arcB* gene. The construct, containing the gentamicin cassette flanked by the upstream and downstream regions of *arcB* was then amplified by PCR with P2266 and P2267, and cloned in plasmid pSW4426T using *EcoRI/PstI* restriction enzymes. To remove the gentamicin cassette, this construct was divergently amplified with primers P2196 and P2197 by PCR on *XhoI* restriction sites, digested with *XhoI* restriction enzyme and ligated. The product, pJGA553, was used to transform *E.coli S17 λpir* by electroporation and confirmed by colony PCR using the primers P1643 and P1728. Plasmid pJGA553 was transferred from *E.coli* to *Vibrio* BB120 by conjugation as mentioned above. The conjugation droplet was streaked in LM supplemented with PB 50 + Cm5 to select for colonies with chromosomal integration of pJGA553. Recombinants were then streaked in LM + arabinose 0.3% (incubate for 24-48hours at 30°C) to induce counter-selection and promote removal of chromosomal *arcB*, generating a clean deletion. Isolated colonies were tested by PCR colony with primers P2186 and P2187 to confirm plasmid excision and *arcB* mutation.

PCR for cloning purposes was performed using the proofreading Bio-X-ACT (Bioline) or Phusion (NEB) enzymes. Other PCRs were performed using Dream Taq polymerase (Fermentas). Digestions were performed with Fast Digest Enzymes (Fermentas), and ligations were performed with T4 DNA ligase (New England Biolabs). All cloning steps were performed in either *E. coli* DH5α, *E. coli S17 λpir* or WT *Ecc15*. All mutants and constructs were confirmed by PCR amplification and subsequent Sanger sequencing performed at the Instituto Gulbenkian de Ciência sequencing facility.

### Construction and selection of *Ecc15* Tn5::kan random insertion mutant library

WT *Ecc15* cells carrying the P*_evf::gfp_* reporter fusion were turned electrocompetent as mentioned above. These electrocompetent cells were transformed with the transposon Tn5::kan (EZ-Tn5™ <KAN-2>Tnp Transposome™ Kit, Lucigen) following the indications of the manufacture. Briefly, 1 µl of Tn5::kan transposome solution was added to 50 µl of WT *Ecc15* electrocompetent cells and using a Bio-rad micropulser (program ECC2) a shock was applied to promote entry of the transposon DNA. Cells were recovered in 1 ml of SOC without shaking for 1 hour, plated in LB + kanamycin (50) and incubated ON for single colonies. Isolated single colonies of the recovered Tn5::kan transformed cells were picked to inoculate 93 wells of a 96 well plate. 2 of the remaining 3 wells were inoculate with the ancestral strain and an empty well was used as negative control. The 96 well plates were grown ON in LB broth supplemented with kanamycin + spectinomycin or LB + spectinomycin (in the case of the ancestral strain), with shaking (700 rpm) at 30°C and frozen at −80°C. For selection of mutants with lower or higher levels of the reporter fusion, each plate was grown for 6 hours in LB + Spec, *evf* reporter fusion was measured using flow cytometry and the mutants with the 10% lower or higher levels in comparison to the WT *Ecc15* strain were isolated to new 96 well plates (masterplates). The masterplates were grown in the same conditions of the first round of selection, and measured 4 independent times. Mutants that were lower or higher than the WT strain in at least 3 out of the 4 measurements were selected and sent to identification of the transposon insertion by whole genome sequencing.

### Tn5::kan insertion identification by whole-genome sequencing

DNA was extracted following a conventional Phenol-Chloroform extraction method. The concentration and purity of DNA were quantified using Qubit and NanoDrop devices, respectively. DNA library construction and sequencing were performed by the IGC genomics facility. Each sample was paired-end sequenced using an Illumina MiSeq Benchtop Sequencer. Standard procedures generated datasets of Illumina paired-end 250 bp read pairs. The reads were filtered using Trimmomatic. Sequences were analyzed using breseq v.0.31.1. An *Ecc15* genome sequenced by the IGC genomic facility using Illumina Miseq and assembly de novo was used as a reference. Insertion sites were identified by aligning the Miseq reads against the *Ecc15* reference genome and the sequence of the Tn5::*kan* transposon. Insertion sites were considered valid when at least 20 sequences containing both a portion of the genome and of the transposon sequence were aligned.

### Plant virulence assay

Plant virulence was analyzed by assessing the maceration of potato tubers with the protocol adapted from (9, 86). Potatoes were washed and surface sterilized by soaking for 10 min in 10% bleach, followed by 10 min in 70% ethanol. Overnight cultures in LB broth were washed twice and diluted to an OD_600_ of 0.05 in phosphate-buffered saline (PBS). Thirty-microliter aliquots were then used to inoculate the previously punctured potatoes. Potato tubers were incubated at 28°C at a relative humidity above 90% for 48 h. After incubation, potatoes were sliced, and macerated tissue was collected and weighed.

### Promoter expression assays

*Ecc15* carrying the different plasmid-borne promoter reporter fusions were grown overnight in LB supplemented with 0,4 % PGA + Spectinomycin (LB PGA+ Spec), inoculated into fresh medium at a starting OD_600_ of 0.05 and incubated at 30°C with aeration. At the indicated timepoints, aliquots were collected to assess growth and the expression of the reporter fusion. For the analyses of reporter expression, aliquots of the cultures were diluted 1:100 in PBS and expression was measured by flow cytometry (LSRFortessa; BD) and analyzed with Flowing Software v 2.5.1, as previously described (87). A minimum of 10,000 green fluorescent protein (GFP)-positive single cells were acquired per sample. Expression of the promoter-*gfp* fusions is reported as the median GFP expression of GFP-positive single cells in arbitrary units. Each experiment included at least 3 independent cultures per genotype, and was repeated on 3 independent days.

### Macro colony and biofilm assays

*Salmonella enterica* serovar Thyphimurium ATCC 14028s strains were grown overnight in LB at 37°C with aeration, subsequently diluted to optical density (OD_600nm_) of 0.05 and incubated for 4 hours at 37°C with aeration in LB. 1 ml of each culture was then washed in 1ml PBS and centrifuged at 13000 r.p.m. for 2min. Washed cultures were resuspended in 1ml PBS, a 5 µl spot was dropped at a center of plates containing LB without salt, which were incubated for 4 days at 28°C. Macro colonies were imaged in a scope with a TCZR036 lens with 0,5x amplification using an mvblue fox3 – 2051ac camera. For biofilm quantification a crystal violet assay was used accordingly to the protocol adapted from (87). Briefly, cultures were grown overnight and diluted to an OD_600_ of 0.05 in 96 well plates containing 100 µl of LB without salt. Plates were incubated statically at 30°C for 90 min to promote adhesion of cell to plastic. Each inoculated well was then washed with 200 µl of PBS twice and filled with 200 µl of fresh media. Cells were left to grow for 24 hours, washed twice with PBS, and stained with 200 µl of 0.1% crystal violet solution for 20 min. The crystal violet was removed by inverting the plate in to a container, washed twice with 250 µl of PBS and left to dry for 30 min. Cell-bound crystal violet was recovered by incubating cells with 200 µl of an 33% glacial acetic acid solution for 15 min. Supernatants were recovered and optical density (OD_580nm_) was measured to quantify biofilm formation.

### Bioluminescence assays

*Vibrio harveyi* strains were grown overnight in AB medium and diluted into fresh AB to an optical density (OD_600nm_) of 0.01. Cells were incubated in a Synergy neo2 plate reader at 30°C with continuous shaking for 8h. Luminescence was determined every 30 min using the luminescence option of the plate reader.

### *Drosophila* Stocks

DrosDel *w^1118^* isogenic stock (*w^1118^ iso*) was used in all experiments (88, 89). Stocks were maintained at 25°C in standard corn meal fly medium composed of 1.1 L water, 45 g molasses, 75 g of sugar, 10 g agar, 70 g cornmeal, 20 g yeast. Food was autoclaved and cooled to 45°C before adding 30 mL of a solution containing 0.2 g of carbendazim (Sigma) and 100 g of methylparaben (Sigma) in 1 L of absolute ethanol. Experiments were performed at 28°C

### Drosophila Infection with *Ecc15*

Egg laying was performed in cages with adult flies at a ratio of three females to one male. To synchronize the egg laying, flies were initially incubated for 1 hour at 25°C, to lay prior fertilized eggs. After this initial incubation, flies were transferred to new cages where eggs were laid for 4 to 6 hours in the presence of standard corn meal fly medium. For bacterial infections, 30 embryos were placed in a 25 ml plastic tube containing 7.5 ml of standard corn meal fly medium with 200 µl of a bacterial cell suspension (OD_600_ = 100 or 1, as specified) from an overnight culture and incubated at 28°C. To assess development and survival of the larvae we counted pupae every 12 hours for 10 days. This allowed us to measure time of development between embryo and pupae, and to calculate percentage of embryos that reached pupal stage (survival).

### Statistical analysis

Statistical analyses were performed in R (90) and graphs were generated using the package ggplot2 (91). All experiments were analyzed using linear mixed-effect models [package lme4, updated version **1.1-20** (92)]. Significance of interactions between factors was tested by comparing models fitting the data with and without the interactions using analysis of variance (ANOVA). Models were simplified when interactions were not significant. Multiple comparisons of the estimates from fitted models were performed with a Tukey HSD (honestly significant difference) test (packages lmerTest (93) and multicomp (94)). To each statistical group a letter is attributed, different letters stand for significant statistical difference.

## Acknowledgments

We thank Joana Amaro for technical assistance, Rita Valente and Roberto Balbontín for critically reading of the manuscript and helpful suggestions. We thank JingTao and Daniel Sobral from the IGC Bioinformatics facility for their help with the analysis of the genomic sequencing.

We acknowledge support from Portuguese national funding agency Fundação para a Ciência e Tecnologia (FCT) for individual grants IF/00831/2015 and IF/00839/2015 to KB.X and L.T., respectively, SRFH/BD/113986/2015 within the scope of the PhD program Molecular Biosciences PD/00133/2012 to F.J.D.V., and project PTDC/BIA-MIC/31984/2017 to L. T..

This work was supported by the research infrastructure ONEIDA projects (LISBOA-01-0145-FEDER-016417 and LISBOA-01-0145-FEDER-022170) co-funded by Fundos Europeus Estruturais e de Investimento from Programa Operacional Regional Lisboa 2020 to K.B.X and L.T.

